# Serotonergic psychedelics delay resting state neurovascular coupling dynamics

**DOI:** 10.64898/2026.10.09.758082

**Authors:** Lautaro Mundel, Vicente Medel, Christopher Timmermann, Patricio Orio, Josefina Cruzat, Carla Pallavicini, Gustavo Deco, Agustin Ibañez, Robin Carhart-Harris, Yonatan Sanz Perl, Enzo Tagliazucchi

## Abstract

Psychedelics are increasingly explored as tools to investigate consciousness and cognition, and as potential treatments for mood disorders. While neuroimaging has been central to characterizing their mechanisms of action, the interpretation of these studies may be confounded by the neurovascular effects elicited by these drugs. Here, we investigated the modulation of resting state neurovascular coupling by the serotonergic psychedelics LSD and DMT. Applying a blind deconvolution algorithm to blood-oxygen-level-dependent (BOLD) signals, we found that both drugs increased the estimated hemodynamic delays in regions overlapping with the distribution of serotonin receptors implicated in psychedelic action and vascular regulation. Deconvolution enhanced the detection of psychedelic-induced changes in whole-brain functional connectivity, suggesting that vascular effects obscure connectivity changes instead of generating spurious results. Finally, a biophysical whole-brain model linked these effects to increased flow-dependent feedback and slower dissipation of the vasoactive drive. These findings underscore the need to account for vascular effects in future psychedelic neuroimaging studies.

## Introduction

Psychedelics are drugs capable of eliciting transient but profound changes in subjective experience, including alterations in perception, mood, and cognition (Aday et al., 2021; Yaden et al., 2021; Cardoso et al., 2023; Moujaes et al., 2025). Recent clinical trials have renewed interest in their therapeutic potential, showing rapid and sustained improvements in conditions such as mood, anxiety and substance abuse disorders (Andersen et al., 2021; Bogenschutz et al., 2022; Davis et al., 2021; Goodwin et al., 2022; Holze et al., 2023; Johnson et al., 2026; Nutt and Carhart-Harris, 2021). Clinical research with psychedelics remains constrained by important methodological challenges, including the difficulty of maintaining effective blinding given their salient subjective effects (Orsini et al., 2026). Although these concerns may be partially addressed through methodological improvements (Szigeti amd Heifets, 2024; Aday et al., 2025), overcoming them will likely require a more comprehensive understanding of the neurobiological mechanisms underlying the therapeutic actions of psychedelics (Muthukumaraswamy, 2023).

In recent years, neuroimaging studies have played a central role in clarifying the neural basis of psychedelic action in the human brain (Erritzoe et al., 2024). Non-invasive magneto- and electroencephalography (M/EEG) studies have identified robust and reproducible biomarkers of the psychedelic state, including broadband reductions in spectral power (Muthukumaraswamy et al., 2013) and increases in entropy and signal complexity (Carhart-Harris et al., 2014; Schartner et al., 2017; Carhart-Harris, 2018). Functional magnetic resonance imaging (fMRI), a technique with comparatively high anatomical resolution, has been widely employed to investigate how psychedelics modulate brain activity and functional connectivity (Girn et al., 2026). To date, the neural correlates of the psychedelic state have been studied primarily with fMRI recordings acquired during resting state, that is, in the absence of explicit task demands or sensory stimulation (Dos Santos et al., 2016; Beneš et al., 2025).

In contrast to M/EEG, fMRI estimates neural activity indirectly by measuring local changes in blood oxygenation associated with the metabolic demands of neural activity (Drew, 2019). As a consequence, the blood-oxygen-level-dependent (BOLD) signal measured with fMRI not only is sensitive to neural activity, but also to vascular parameters such as vessel diameter and stiffness, as well as to the coupling between neural activity and the associated vascular response, known as neurovascular coupling (NVC) (Hillman, 2014). Because of this dependence on non-neural factors, fMRI faces important interpretability challenges when used to study pharmacological conditions that affect the cardiovascular system, such as the acute effects induced by serotonergic psychedelics (Cole et al., 2010).

Psychedelics act primarily as partial agonists at the 5-HT2A receptor (Nichols, 2016; Kwan et al., 2022), which together with other serotonin receptor subtypes, is implicated in the regulation of blood vessel diameter (Berger et al., 2009). In particular, the agonist or partial agonist activity of psychedelics at serotonin receptors may influence vascular tone, as these receptors are expressed in vascular smooth muscle and endothelium, where their activation can produce vasoconstriction or vasodilation (Wsół, 2023). Recent animal optical imaging studies have demonstrated that psychedelics disrupt the NVC during task and rest conditions, decoupling neural activity from changes in cerebral blood flow, and increasing the latency of the hemodynamic response while also shortening its duration (Padawer-Curry et al., 2025; Zirkel et al, 2026). In humans, psychedelics can alter task-elicited hemodynamic responses although their neurovascular effects during the resting state remain to be established (Padawer-Curry et al., 2025).

Here, we investigated the modulation of resting state NVC in humans by two serotonergic psychedelics, dimethyltryptamine (DMT) and lysergic acid diethylamide (LSD). Across the whole brain, we implemented a BOLD signal blind deconvolution algorithm to compute a proxy of the hemodynamic response function (HRF) to spontaneous fluctuations in neural activity (Wu et al., 2013). This procedure allowed us to estimate key resting state HRF shape parameters—height, width, and latency—compare them between drug and placebo conditions, and contrast the anatomical distribution of psychedelic-induced alterations with the cortical density of serotonin receptors (Hansen et al., 2022). Using the deconvolved BOLD signals, we then investigated whether the estimated vascular effects could lead to false positives or false negatives in analyses of psychedelic-induced changes in whole-brain functional connectivity and global signal variance. Finally, we fitted a biophysical whole-brain model to reproduce functional connectivity under drug and placebo conditions, with the objective of disentangling neural and vascular contributions, and informing the potential mechanisms underlying the effects of psychedelics on the dynamics of vasodilatory signals (Friston et al., 2000; Deco et al., 2018).

Based on previous animal studies, we expected LSD and DMT to increase the latency of the resting state hemodynamic response, as reflected in the HRF time-to-peak parameter (Padawer-Curry et al., 2025; Zirkel et al, 2026). We further hypothesized that vascular effects related to vasoconstriction would occur in regions overlapping with the density of 5-HT2A and 5-HT1B receptors, but not with the distribution of 5-HT1A receptors, which are primarily associated with vasodilation (Berger et al., 2009). After BOLD signal deconvolution, we expected changes in whole-brain FC and global signal variance to reveal false positives or false negatives associated with vascular confounds (Tak et al., 2015). Finally, we hypothesized that LSD and DMT would induce consistent alterations in biophysical parameters related to the regulation of the NVC, consistently with a delayed HRF peak.

## Results

### Methodological overview

A methodological overview is provided in Fig. 1. We analyzed pre-existing datasets of resting state fMRI recordings acquired from participants under the acute effects of two serotonergic psychedelics: LSD (75 μg; N = 15; four female, mean age 30.5 ± 8.0 years; Carhart-Harris et al., 2016) and DMT (20 mg; N = 17; 5 females, mean age 39.6 ± 9.6 years; Timmermann et al., 2023) (Fig. 1, “Data acquisition”). In both datasets, eyes-closed resting state fMRI data were also acquired after participants received an inactive placebo consisting of saline solution, following a blinded (double-blind for LSD, single-blind for DMT), randomized, crossover and counterbalanced design. Further experimental details, including fMRI data acquisition and preprocessing procedures, are provided in the Methods section and the supplementary material, sections “Experimental procedure (LSD)” and “Experimental procedure (DMT)”.

**Figure 1.**
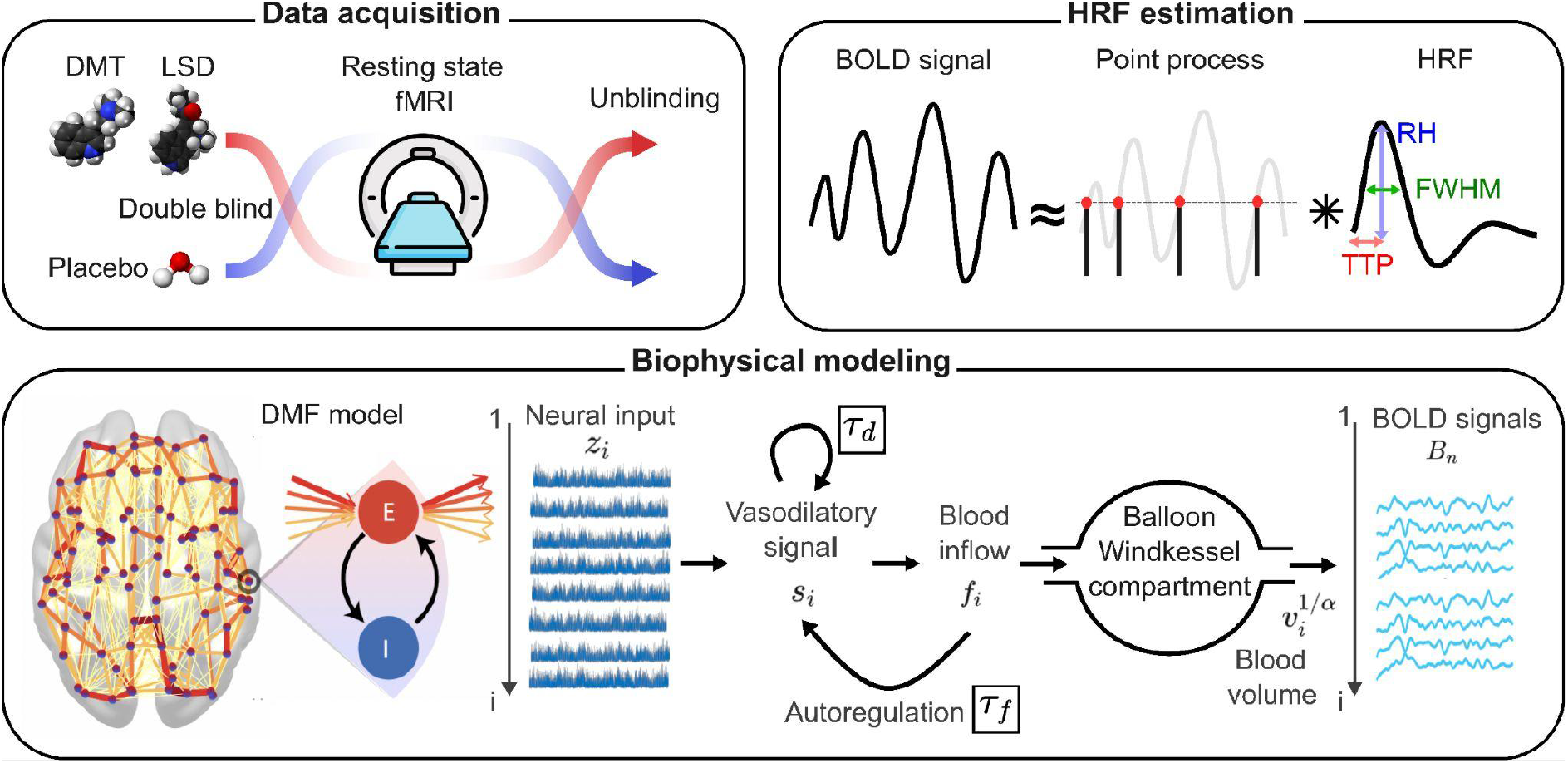
Methodological overview. Data acquisition followed a double-blind (LSD) or single-blind (DMT), placebo-controlled, crossover and counterbalanced design, with resting state fMRI data obtained from human participants under the acute effects of LSD (75 μg) or DMT (20 mg) and under a control condition consisting of saline solution. HRF estimation was performed using a BOLD deconvolution algorithm, which identified a point process underlying large-amplitude BOLD events and used it as input to a Wiener deconvolution algorithm, yielding an estimate of the convolution kernel (i.e. the resting state HRF), as well as a deconvolved version of the original BOLD time series. Biophysical modelling was based on the dynamic mean-field model (DMF), consisting of local excitatory (E) and inhibitory (I) neural populations, with long-range E–E interactions determined by the human connectome. The simulated excitatory firing-rate time series (z_i_) were converted into BOLD signals using the Balloon–Windkessel model, with time constants τ_*d*_ and τ_*f*_ for the spontaneous vasodilatory signal decay and the flow-dependent autoregulatory feedback, respectively.

In fMRI recordings, the HRF is typically estimated using active tasks or stimulation paradigms (Lindquist et al., 2009) or through interventions designed to induce a vascular response decoupled from neural activity, such as a hypercapnic challenge (Chang et al., 2008). However, these methods require active participation from the subjects and thus are unable to estimate the HRF associated with spontaneous neural activity fluctuations. To overcome this limitation, we implemented a resting state BOLD signal blind deconvolution procedure (Fig. 1, “HRF estimation”) based on the detection of spontaneous large-amplitude BOLD events (Wu et al., 2013; Wu et al., 2021). The timing of these events was encoded as a point process, approximating a discrete set of neural events driving the increases in the BOLD signal (Tagliazucchi et al., 2012). The resting state HRF was estimated from the BOLD response to these events and then used to deconvolve the BOLD signal time series (Wu et al., 2013). Finally, we extracted three shape parameters from the estimated HRF: response height (HR), full-width at half maximum (FWHM), and time to peak (TTP), as illustrated in Fig. 1 (“HRF estimation”, right panel).

To investigate the dynamical mechanisms underlying psychedelic-induced alterations to the HRF, we used a generative biophysical model of whole-brain activity (Fig. 1, “Biophysical modelling”). The dynamic mean-field model (DMF; Wong and Wang, 2006; Deco et al., 2018) comprises local excitatory (E) and inhibitory (I) neural populations, with local E–I interactions and long-range E–E interactions constrained by the anatomical connectome, i.e. the network of direct physical connections between brain regions estimated using diffusion tensor imaging (DTI) and scaled by the global parameter *G* (Sporns et al., 2011). At the i-th region of interest (ROI), the simulated excitatory firing-rate time series (z_i_) drives a vasodilatory signal (s_i_) leading to changes in the local blood inflow (f_i_) (Stephan et al., 2007). This process is governed by the spontaneous decay of s_i_ with time constant τ_*d*_, and by flow-dependent autoregulatory feedback with time constant τ_*f*_. Blood volume (v_i_) is then determined from blood inflow (f_i_) using a Balloon–Windkessel model (Friston et al., 2000). The model therefore produces a simulated BOLD signal in each region, whose properties can be compared with the empirical signal to optimize the phenomenological neurovascular response time constants τ_*d*_ and τ_*f*_.

### Psychedelic-induced changes to resting state HRF latency

We first compared the estimated HRF shape parameters between each drug condition and its corresponding placebo condition (PCB). To reduce the number of comparisons and improve interpretability, ROIs were grouped into major resting state networks (RSNs), comprising cortical and subcortical regions with coordinated spontaneous activity fluctuations and shared functional roles (Beckmann et al., 2005). These RNSs included primary visual (Vis), extrastriate visual (ES), auditory (Aud), sensorimotor (SM), default mode (DM), and executive control (EC) networks. Further information on the anatomical regions included in each RSNs is provided in the supplementary material (section “Resting state networks”).

Figure 2A shows the estimated values of the time-to-peak (TTP) parameter, which ranged between 4 and 7 s, consistent with previous studies of resting state and task-evoked hemodynamic responses (Lindquist et al., 2009; Wu et al., 2013). Comparisons between drug and placebo conditions revealed widespread psychedelic-induced increases in hemodynamic response latency. Both LSD and DMT increased TTP in primary and extrastriate visual regions, while DMT additionally increased TTP in all RSNs except the DM network. Comparisons of the remaining HRF shape parameters (RH and FWHM) did not reveal significant differences between drug and placebo conditions (supplementary figure S1).

**Figure 2.**
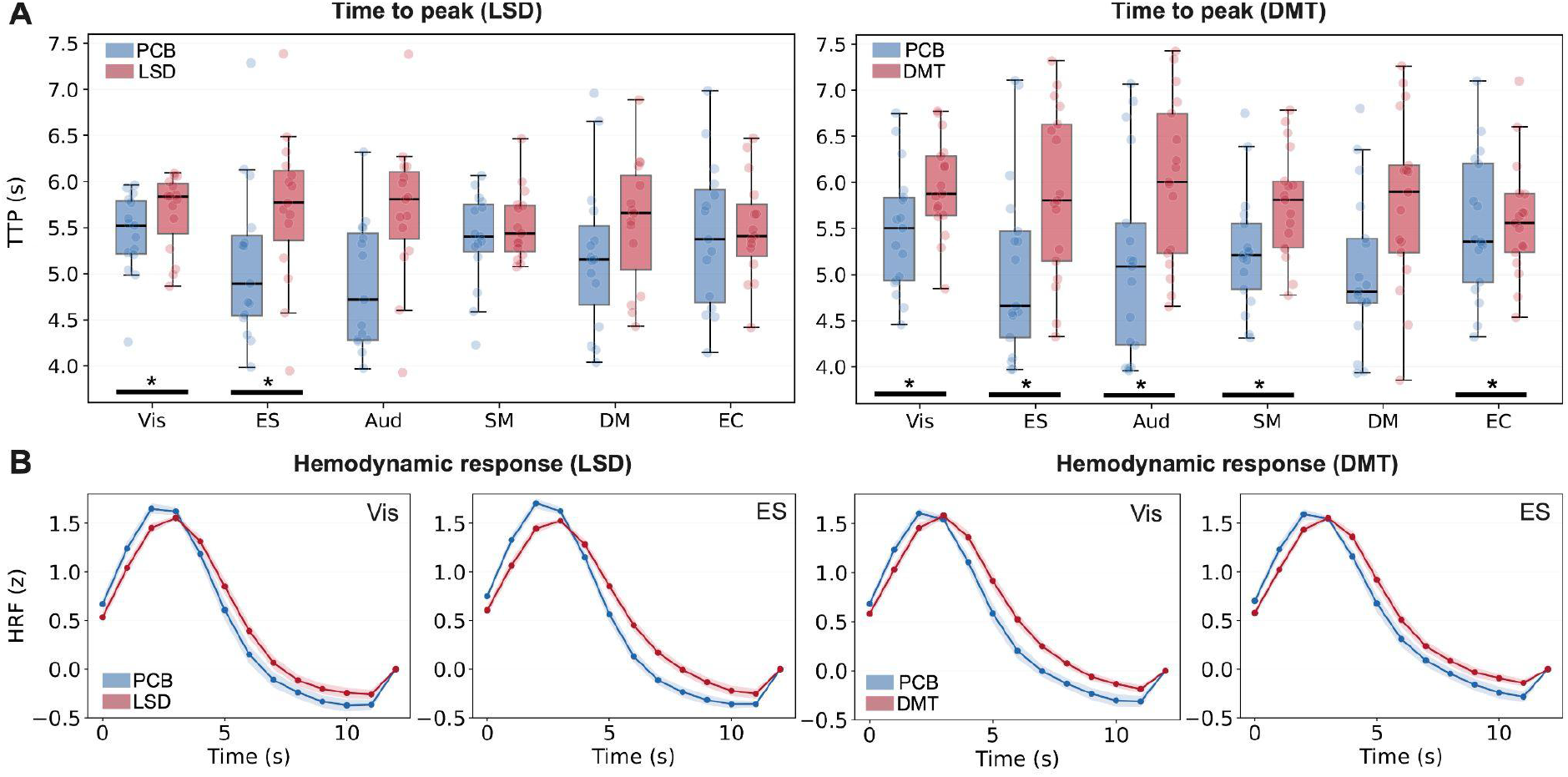
LSD and DMT induce widespread increases in resting state HRF latency. **A)** Boxplots of the estimated HRF time to peak shape parameter (TTP) for LSD and DMT relative to placebo (PCB), shown separately for six major resting state networks (RSNs): primary visual (Vis), extrastriate visual (ES), auditory (Aud), sensorimotor (SM), default mode (DM), and executive control (EC). Asterisks indicate significant drug–placebo differences (p < 0.05, FDR-corrected for multiple comparisons). **B)** Estimated HRFs computed at Vis and ES networks, illustrating the delayed vascular response during the acute effects of LSD and DMT.

The increased latency of the hemodynamic response is illustrated in Fig. 2B. This panel presents the average resting state HRFs computed at two RSNs where TTP differences were observed for both drug conditions relative to placebo (ES and VIS networks). The estimated HRFs followed a shape consistent with previous descriptions of vascular responses to the metabolic demands of neural activity. As shown in Fig. 2B, psychedelics increased the latency of this response without markedly distorting its overall shape, resulting in a delayed peak and a more sustained response after 4 s.

### Overlap between TTP increases and receptor density

We next investigated the anatomical distribution of psychedelic-induced increases in TTP. Figure 3A shows ROIs with t > 1.5 for the comparison between each drug condition and placebo (paired Student t-test). Although none of these ROIs reached the threshold for statistical significance after correction for multiple comparisons (α=0.05), the spatial pattern of TTP changes was similar for LSD and DMT. In particular, TTP increases were predominantly located in frontal, occipital, and inferior temporal regions, as well as in anterior and central midline regions.

**Figure 3.**
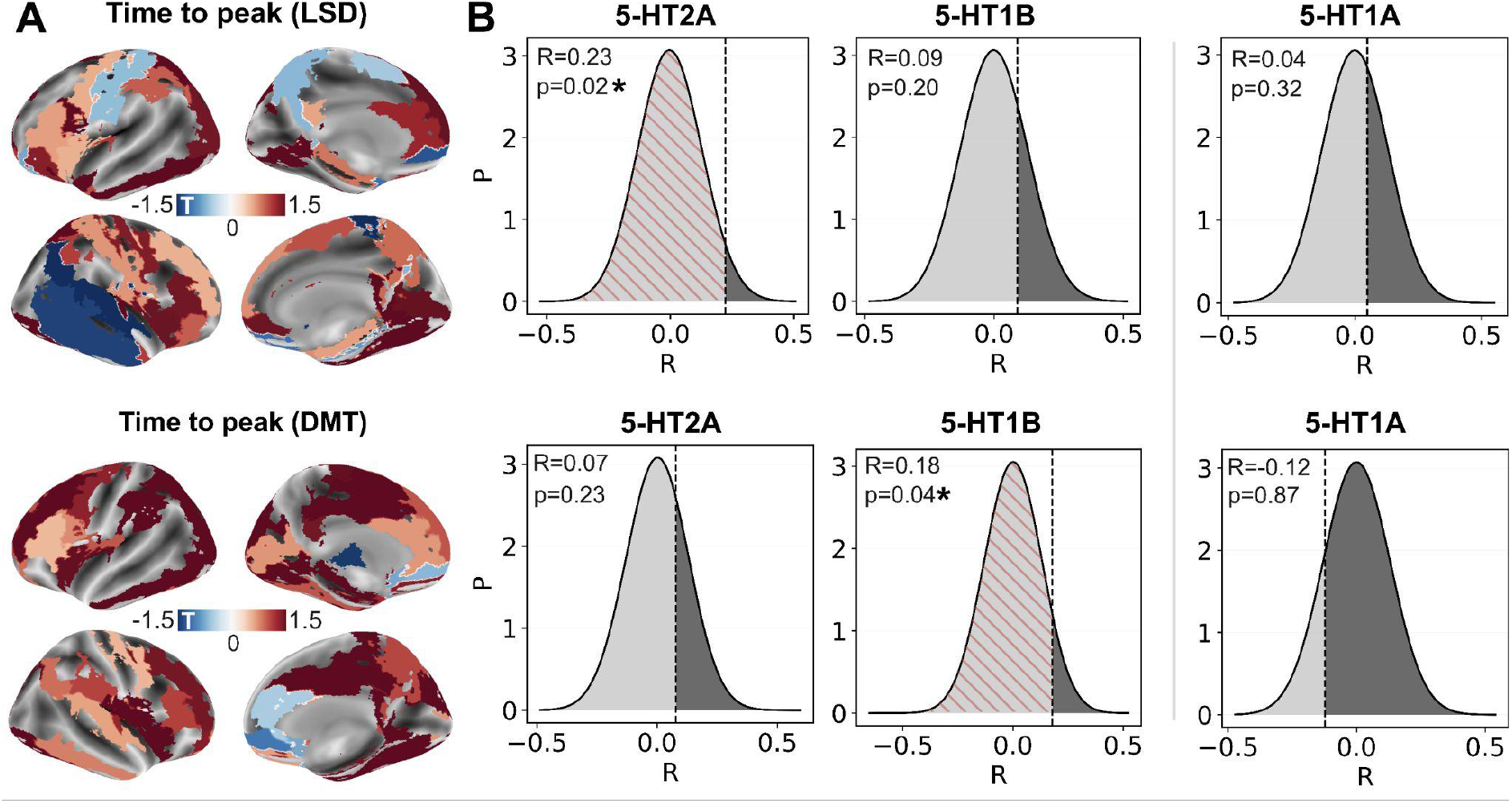
The anatomical distribution of TTP increases correlates with the density of 5-HT2A and 5-HT1B serotonin receptor subtypes. **A)** Rendering of regional t-values for the drug–placebo comparison, obtained using two-tailed t-tests and thresholded at t > 1.5. Results are shown for LSD and DMT in the top and bottom panels, respectively. **B)** Pearson correlation coefficients (R) and corresponding p-values for the associations between regional t-values and receptor density values measured with PET. Histograms show the null distributions of correlation coefficients obtained from 50,000 independent permutations of receptor density values. Dark and light gray bars indicate permutations yielding correlation coefficients larger and smaller than the empirical value, respectively. Asterisks indicate significant correlations (p < 0.05, FDR-corrected).

Figure 3B compares these spatial patterns with the cortical distribution of three serotonin receptor subtypes: 5-HT2A, 5-HT1B, and 5-HT1A, shown in the first, second, and third columns, respectively. Among the PET maps published by Hansen et al. (2022), these receptors were selected because of their involvement in vasoregulation and their distinct profiles of vasoconstrictive and vasodilatory effects. Specifically, the vascular effects of receptors 5-HT2A and 5-HT1B are primarily linked to vasoconstriction, while agonism of the 5-HT1A receptor is associated with vasodilatory effects (Berger et al., 2009). The inset in each panel shows the Pearson correlation coefficient (R) and corresponding p-value for the association between regional t-values from the drug–placebo comparison and regional receptor density, as measured in previous positron emission tomography (PET) studies (Hansen et al., 2022). Statistical significance was assessed using a permutation test with 50,000 independent random permutations of regional receptor density values; the resulting null distributions of correlation coefficients are shown in the histograms.

This analysis revealed significant positive correlations between regional t-values and cortical receptor density for 5-HT2A receptors under LSD (R = 0.23) and 5-HT1B receptors under DMT (R = 0.18) (Fig. 3B). In contrast, the anatomical distribution of 5-HT1A receptors was not correlated with regional t-values for either compound (Fig. 3B, last column).

### Functional connectivity and BOLD signal deconvolution

To investigate the impact of psychedelic-induced vascular effects on estimates of functional connectivity, we compared FC between RSNs in drug and placebo conditions before and after BOLD signal deconvolution. Figure 4A (“BOLD FC”) shows FC changes for LSD versus PCB (top) and DMT versus PCB (bottom). Although no differences survived correction for multiple comparisons, both drugs presented widespread trends toward reduced coupling of spontaneous BOLD signal fluctuations involving the SM, Vis, and ES networks. For DMT, we also observed a tendency toward increased FC between the SM and Aud networks, and between the EC network and both the SM and Aud networks.

**Figure 4.**
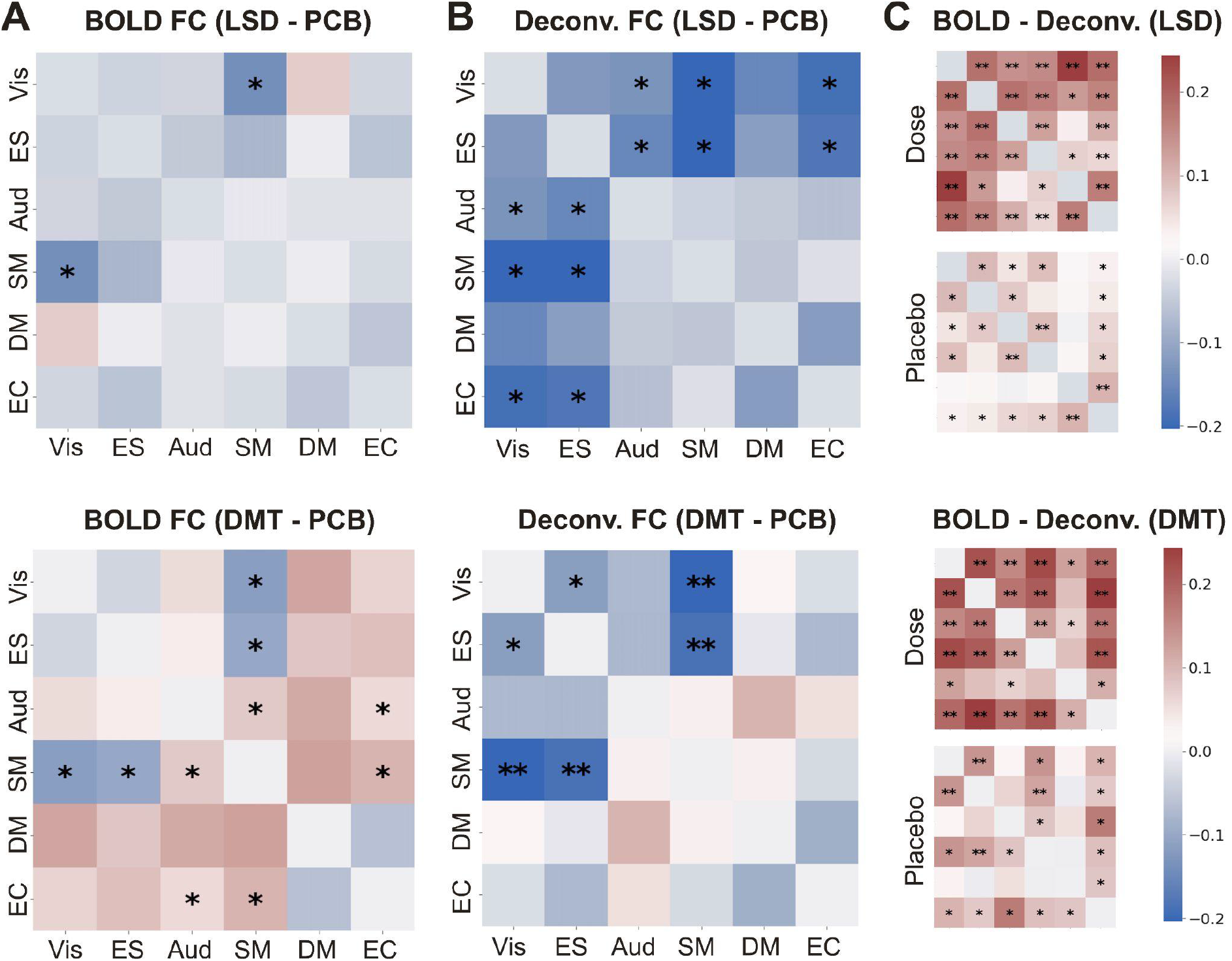
Deconvolution enhances LSD- and DMT-induced BOLD signal desynchronization. **A)** Drug–placebo differences in functional connectivity (FC) matrices for LSD and DMT, computed from the original BOLD signals. **B)** Drug–placebo differences in FC matrices computed from the deconvolved BOLD signals. **C)** Differences in FC matrices between original and deconvolved BOLD signals, shown separately for drug and placebo conditions. Color bars indicate t-statistics from two-tailed t-tests. Asterisks indicate uncorrected significance levels (*p < 0.05, **p < 0.001) (Vis: primary visual, ES: extrastriate visual, Aud: auditory, SM: sensorimotor, DM: default mode, EC: executive control).

FC reductions became more widespread and pronounced after deconvolution (Fig. 4B, “Deconv. FC”), although they did not reach corrected statistical significance. Under LSD, deconvolution revealed reduced FC among the EC, Vis, and ES networks, as well as additional reductions between the SM and ES networks, and between the Aud network and both visual networks. Under DMT, deconvolution increased the magnitude of FC reductions already present in the original BOLD signals and additionally revealed reduced connectivity between the Vis and ES networks. In contrast to LSD, deconvolution under DMT also eliminated the uncorrected FC increases observed in the original BOLD signals.

Comparing FC matrices computed from original and deconvolved signals showed that the net effect of deconvolution was to decrease FC between RSNs (Fig. 4C, “BOLD ™ Deconv.”). This reduction was more prominent in the drug conditions, suggesting a contribution of psychedelic-induced vascular effects. Because deconvolution selectively reduced FC estimates in the drug conditions, its net effect was to accentuate the FC reductions already observed in the original BOLD time series.

### Global signal variance and BOLD signal deconvolution

Next, we addressed the effect of deconvolution on fluctuations in the global resting state BOLD signal. In particular, we investigated whether this procedure could change drug vs. placebo differences in global signal variance. As shown in the supplementary figure S2, LSD reduced global BOLD signal variance relative to placebo, whereas this effect was not observed under DMT. We also found that deconvolution reduced global signal variance for both LSD and DMT. In the case of LSD, the drug–placebo difference in global signal variance was no longer significant after deconvolution, suggesting that the apparent reduction observed in the original BOLD signal was influenced by vascular effects.

### Biophysical modelling of whole-brain activity and neurovascular coupling

We employed the DMF model to disentangle neural and vascular contributions to drug-induced changes in resting state BOLD signals. Following previous studies (Deco et al., 2019; Kringelbach et al., 2020), we first optimized the global coupling parameter, *G*, which scales the inter-areal structural connectivity matrix, or connectome. Optimization was performed by maximizing the structural similarity index measure (SSIM) between empirical and simulated FC matrices, which represents an intermediate measure between Euclidean and correlation similarities (Sanz Perl et al., 2021). Figure 5A shows SSIM values as a function of *G* for LSD and DMT. In both datasets, the optimal value of *G* was lower in the drug condition than in the corresponding placebo condition, consistent with previous reports of psychedelic-induced BOLD signal desynchronization (Siegel et al., 2025; Girn et al., 2026).

**Figure 5.**
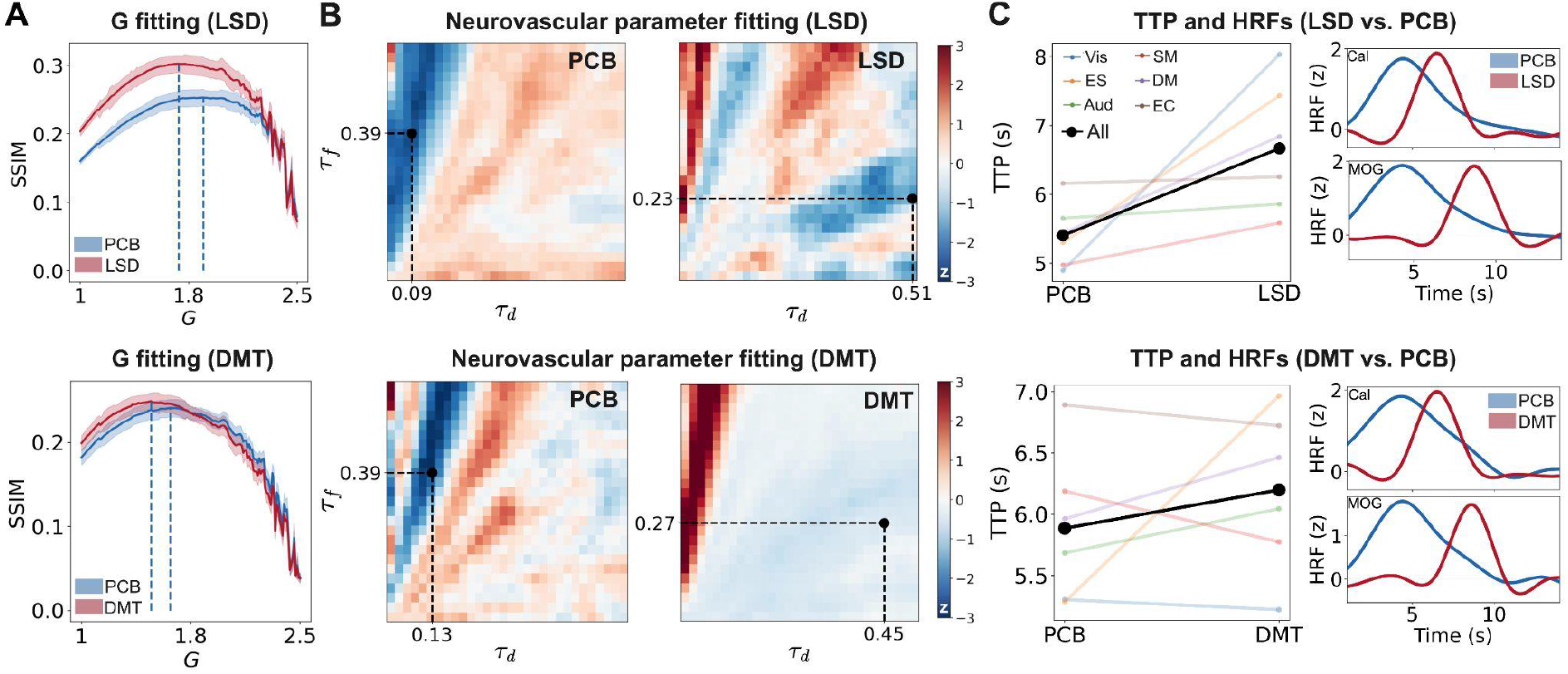
Increased TTP under psychedelics is associated with changes in vasodilatory signal decay and flow-dependent autoregulation. **A)** Optimization of the connectome scaling parameter *G*, quantified using the structural similarity index measure (SSIM) between empirical and simulated FC matrices, shown for LSD and DMT together with their corresponding placebo (PCB) conditions. **B)** Difference between empirical and simulated global HRF TTP values, expressed as z-scores, as a function of neurovascular coupling time constants τ_*d*_ and τ_*f*_. Dashed lines indicate the optimal parameter combination. **C)** Left panels: TTP values of simulated BOLD time series for the six RSNs included in this study, contrasting LSD and DMT with their corresponding placebo conditions. Right panels: representative simulated HRFs for ROIs located in primary visual cortex, the calcarine sulcus (Cal), and medial visual cortex, the middle occipital gyrus (MOG), shown for LSD and DMT together with HRFs estimated from the placebo conditions (Vis: primary visual, ES: extrastriate visual, Aud: auditory, SM: sensorimotor, DM: default mode, EC: executive control).

After setting the optimal value of parameter *G*, we determined the NVC phenomenological model parameters that best reproduced the psychedelic-induced increases in hemodynamic response latency. Figure 5B shows the results of systematically varying the global value of parameters τ_*d*_ and τ_*f*_, corresponding to the time constants for the self-decay of the vasodilatory signal and the flow-dependent autoregulatory feedback, respectively. For each combination of τ_*d*_ and τ_*f*_, we applied the resting state deconvolution procedure to the simulated BOLD signals and compared empirical and simulated HRF latencies.

The difference between empirical and simulated TTP values showed distinct configurations for drug and placebo conditions. Time constants τ_*d*_ and τ_*f*_ were not independent, as evidenced by the regions of parameter space yielding the optimal SSIM values. Consistent patterns were observed for both psychedelic drugs and their corresponding placebo conditions. For both LSD and DMT, the optimal drug-condition parameters were τ_*d*_ ≈ 0.2 and τ_*f*_ ≈ 0.5. In the placebo conditions of both experiments, τ_*f*_ was larger (τ_*f*_ ≈ 0.4) and τ_*d*_ was lower (τ_*d*_ ≈ 0.1). Notably, while changes to parameters τ_*d*_ and τ_*f*_ modulated the HRF, the similarity between empirical and simulated FC remained comparatively stable (supplementary figure S3), supporting the decoupling of vascular and neural effects under psychedelics.

As expected, these changes in neurovascular coupling parameters increased resting state HRF latency across multiple RSNs (Fig. 5C). For LSD, this increase was observed across all networks, whereas for DMT it was observed in all networks except VIS, EC, and SM. The right panels of Fig. 5C show representative simulated HRFs for ROIs in primary visual cortex, the calcarine sulcus (Cal), and medial visual cortex, the middle occipital gyrus (MOG). Similar to the empirical data, the simulated HRFs were shifted in time, resulting in larger TTP values. We also observed a tendency toward broader HRF responses under placebo, however, this was not evident in the empirical HRF estimates.

## Discussion

Ever since its discovery in 1935, serotonin has been implicated in the regulation of vascular tone (Berger et al., 2009). The vasoregulatory role of serotonin is reflected in its etymology, derived from *serum* (blood serum) and the Greek *tonos* (tension or stretching). The modern history of psychedelics also highlights the close interactions between these drugs and the cardiovascular system. Perhaps the best-known example is the discovery of LSD in 1938 by Albert Hofmann, who at the time was investigating vasoconstrictive ergot alkaloids for the treatment of postpartum hemorrhage (Hofmann, 1980). As early as 1954, it was suggested that psychedelics might exert their physiological effects by antagonizing serotonin action in the human brain (Gaddum & Hameed, 1954). This proposal was later revised by a series of experiments showing that partial agonism at the 5-HT2A receptor is the main pharmacological mechanism underlying psychedelic action (Geyer, 2024).

Neuroscientific and clinical research with psychedelics has largely focused on their distinctive subjective effects and their neural correlates, while paying comparatively less attention to their impact on human physiology. However, the widespread use of modern neuroimaging technologies to study these compounds has renewed interest in their cardiovascular effects, especially those involving changes in the NVC. The metabolic demands associated with neural activity trigger a complex cascade of processes that produces local changes in cerebral blood flow and oxygenation, which contribute to the BOLD signal measured with fMRI (Logothetis and Wandell, 2004; Drew, 2019). However, the BOLD signal does not depend solely on oxygen metabolism; it is also sensitive to vascular parameters and to alterations in NVC associated with oxidative stress, inflammation, endothelial dysfunction, altered neuron–astrocyte signaling, and other processes (Petzold and Murthy, 2011; Guerra et al., 2018; Uhlirova et al., 2016; Rosenegger and Gordon, 2015; Schaeffer and Iadecola, 2015). As a result, BOLD signals can be confounded in conditions such as Alzheimer’s disease and stroke, which are characterized by compromised vascular integrity and impaired signaling among neurons, astrocytes, and blood vessels (Girouard and Iadecola, 1985; Zhu et al., 2022).

Mounting evidence suggests that psychedelics can introduce vascular confounds into BOLD signal time series and derived functional connectivity measures. Two recent animal studies have shown that hallucinogenic 5-HT2A agonists can decouple neural activity from hemodynamics, both during resting state and in response to sensory stimulation. Using two-photon imaging combined with wide-field multimodal imaging, Zirkel and colleagues investigated how psilocybin modulates stimulus-induced neurovascular responses in the visual cortex (Zirkel et al., 2026). They found that psilocybin prolonged the blood-flow response to stimulation without altering vessel diameter or blood-flow velocity, suggesting an effect specific to NVC. Specifically, this prolongation reflected a more sustained hemodynamic response, without significant changes in the TTP parameter.

An independent study by Padawar-Curry et al. administered 2,5-dimethoxy-4-iodoamphetamine (DOI) to awake mice while monitoring ensemble-level neural activity with wide-field optical imaging (Padawar-Curry et al., 2025). In agreement with other studies, hallucinogenic doses of DOI decoupled neural activity from the associated hemodynamic response. Although the temporal properties of stimulus-evoked neural activity were not modulated by DOI, the hemodynamic response decreased after DOI and exhibited a more negative post-stimulus undershoot, consistent with changes in stimulus-evoked NVC. Importantly, DOI-induced modulation of resting state NVC showed key differences from those reported by Zirkel and colleagues, including increased TTP and decreased FWHM (Zirkel et al., 2026). These discrepancies may reflect drug-specific effects, as well as state-dependent modulation of NVC by psychedelics, with delayed TTP occurring in response to spontaneous neural activity fluctuations, but not necessarily in response to sensory stimulation.

Our results from human resting state fMRI are partially aligned with these findings, with some discrepancies that may reflect differences between task-evoked and resting state conditions. DOI narrowed the stimulus-evoked HRF (Padawar-Curry et al., 2025), contrasting with our finding of preserved FWHM under both LSD and DMT at rest. We found that LSD and DMT increased the TTP of the estimated HRF, in agreement with the increased resting state hemodynamic latency under DOI reported by Padawar-Curry et al. Although Zirkel et al. did not report changes in TTP, their observation of a more sustained hemodynamic response under psilocybin is consistent with our HRF estimates, as exemplified in Fig. 2B (Zirkel et al., 2026). Interestingly, Padawar-Curry et al. also analyzed data from a previously published human fMRI study of psilocybin, showing that the drug decreased the TTP of the task-evoked HRF (Padawar-Curry et al., 2025), further supporting the possibility that psychedelics modulate neurovascular coupling differently during task and rest.

The psychoactive effects of psychedelics depend primarily on agonism at the 5-HT2A receptor (Nichols, 2016; Kwan et al., 2022). In animal studies, pretreatment with 5-HT2A antagonists reduced the effects of DOI and psilocybin on NVC, suggesting that this receptor is also implicated in psychedelic modulation of the HRF (Padawar-Curry et al., 2025; Zirkel et al., 2026). Activation of 5-HT2A receptors in vascular smooth muscle cells can trigger vasoconstriction (Nishihira et al., 2008; Berger et al., 2009); however, psychedelics also show significant affinity for other serotonin receptor subtypes (Ray, 2010; Rickli et al., 2016) that are associated with opposing effects on vascular tone, such as the 5-HT1A receptor (Kakoki et al., 1999; Blessing et al., 2004). Leveraging PET receptor-density maps and the anatomical coverage of whole-brain fMRI (Hansen et al., 2022), we investigated correlations between psychedelic-induced changes in HRF TTP and the cortical distribution of 5-HT1A, 5-HT1B, and 5-HT2A receptors. This analysis revealed significant associations between HRF latency and the density of 5-HT2A and 5-HT1B receptors under the acute effects of LSD and DMT, respectively. Together with the absence of significant correlations with 5-HT1A receptor density, these findings suggest that the vasoconstrictive effects of psychedelics may contribute to resting state HRF alterations. However, this mechanism cannot be unequivocally inferred from our analysis, given that both LSD and DMT could exert complex vasoregulatory effects due to their interactions at other serotonin receptor subtypes.

Psychedelics elicit heterogeneous and scale-dependent effects on the functional connectivity of resting state BOLD signals (Yu et al., 2024; Farré-Colomés et al., 2026). Recently, psilocybin has been implicated in the massive disruption of cortical and subcortical resting state FC (Siegel et al., 2025), while a meta-analysis integrated 11 resting state studies showed desynchronization between specific RSNs, including visual, sensorimotor and limbic networks (Girn et al., 2026). In agreement with these reports, we observed that both LSD and DMT reduced inter-network connectivity, although these effects reached only an uncorrected threshold for statistical significance. Importantly, these FC reductions were amplified after deconvolving resting state BOLD signals with the estimated HRF kernel, resulting in stronger and more widespread FC decreases relative to placebo. In the case of DMT, increased FC between the EC, Aud, and SM networks disappeared after deconvolution. With these exceptions, the vascular confounds estimated from the resting state HRF tended to produce false negatives by obscuring psychedelic-induced FC changes, rather than exaggerating FC differences relative to placebo. This possibility is consistent with previous animal studies and may reflect heterogeneous changes in TTP across the cortex (Fig. 3A) (Padawar-Curry et al., 2025; Zirkel et al., 2026), which could introduce spurious phase differences between time series and thereby disrupt FC estimates in the drug condition.

In the temporal domain, signal deconvolution eliminated differences in global signal variance that were observed for LSD relative to placebo. The global signal has been associated with spontaneous fluctuations in neural activity (Schölvinck et al., 2010), and changes in its amplitude or variance have been reported across brain states characterized by altered levels of vigilance (Fukunaga et al., 20026; Song et al., 2022), as well as altered conscious content, including the acute effects of serotonergic psychedelics (Tagliazucchi et al., 2014). Deconvolution may introduce a form of normalization, whereby differences in signal amplitude are compensated by the response height (RH) parameter of the estimated HRF kernel. However, because our data-driven procedure estimated this parameter from the signal itself, this analysis entails a degree of circularity. Future studies should estimate the HRF through independent approaches, such as stimulation or task paradigms, concurrent electrophysiological recordings, or other external measures of neural activity, and evaluate how deconvolution with independently estimated kernels affects the amplitude and variance of BOLD signals.

The latency of the HRF could increase if vasodilatory signaling is slowed, for example through altered decay of fast-turnover messengers such as nitric oxide, attenuated conducted vasodilation along electrically coupled vascular cells, or changes in feedback-controlled arteriolar compliance and vascular mechanics (Behzadi and Liu, 2005; Jensen and Holstein-Rathlou, 2013; Hosford and Gourine, 2019). Another potentially contributing factor is delayed negative feedback within the neurovascular response. Activity-evoked increases in flow may wash out vasoactive metabolites, while restoration of tissue oxygenation may reduce hypoxia-sensitive vasodilation (MacVicar and Newman, 2015; Nippert et al., 2018; Hosford and Gourine, 2019). In parallel, myogenic autoregulation, astrocyte-derived vasoactive signaling, and neurogenic modulation of cerebrovascular tone could shape the recovery dynamics and temporal dispersion of the HRF (Hamel et al., 2006; Koehler and Roman, 2009; Koller and Toth, 2012). Resting state fMRI is inherently limited in its ability to identify the precise neurovascular processes underlying the observed changes in HRF TTP. However, implementing a whole-brain dynamic mean-field model with neurovascular coupling described by a Balloon–Windkessel model allowed us to estimate the relative contributions of vasodilatory self-decay and flow-dependent autoregulation, with the characteristic time of these processes encoded in model parameters τ_*d*_ and τ_*f*_, respectively. While these phenomenological time constants are not associated with a unique biophysical mechanism, they represent distinct contributions to the dynamics of the vasodilatory signal which can be selectively modified by psychedelics.

Our results indicate that LSD and DMT simultaneously slow down the initial vasoactive signal, reflected in larger τ_*d*_ values, while strengthening feedback control of blood inflow, reflected in increased τ_*f*_ (Stephan et al., 2007). Although these changes are opposing, their combined effect could delay the HRF if the prolonged vasoactive drive overcompensates for the tighter coupling with blood inflow. The increase in τ_*d*_ suggests that psychedelics lengthen the lifetime of the upstream vasodilatory signal, potentially through direct serotonergic modulation of astrocytic, endothelial, or vascular smooth-muscle pathways (Ishida et al., 1998; Cohen et al., 1999; Zhang et al., 2010; Machida et al., 2013). Conversely, larger values of time constant τ_*f*_ indicate increased effective autoregulatory feedback on blood inflow (Stephan et al., 2007). Given the known vasoactive properties of serotonergic psychedelics, including serotonin-mediated effects on vascular smooth muscle (Wsół, 2023; Nahlawi et al., 2025), one possible interpretation is that drug-induced vasoconstriction shifts the vascular operating point and modifies myogenic or flow-dependent feedback mechanisms, although further studies leveraging other imaging methodologies are necessary to clarify the underlying neurobiological mechanisms.

In agreement with previous empirical findings and modelling efforts, our results show that vascular changes may unfold independently of the neural effects induced by psychedelics (Padawar-Curry et al., 2025; Zirkel et al., 2026). After optimizing the coupling parameter G for each drug condition and its corresponding placebo condition, optimization of the neurovascular coupling parameters did not significantly alter the model fit to empirical FC. Thus, the same generative model of regional neural activity could produce distinct hemodynamic profiles depending on the regulation of the vasodilatory signal. This decoupling between neural and vascular responses to psychedelics represents an important concern for future neuroimaging studies and should be further investigated by integrating animal experiments, human neuroimaging, and computational modelling.

Our study has some important limitations. The most salient is the lack of independent validation of the resting state HRF estimates, such as that provided by simultaneous electrophysiological recordings. Previous studies have related psychedelic-induced changes in hemodynamic activity to the onset of sensory stimulation or task execution; however, these approaches are not well suited to address the vascular effects of psychedelics during resting state (Chang et al., 2008; Lindquist et al., 2009; Prokopiou et al., 2022). Simultaneous EEG–fMRI recordings of spontaneous brain activity could allow estimation of the transfer function between neuroelectrical and hemodynamic measures (Lu et al., 2006; de Munck et al., 2007), as well as how this function is altered during the acute effects of psychedelics (Timmermann et al., 2023). Combined with biophysical modelling to investigate the underlying mechanisms, this approach could help disentangle vascular confounds in fMRI time series acquired during acute psychedelic states.

In summary, we provided evidence that psychedelics alter neurovascular coupling during resting state, increasing the latency of the hemodynamic response in networks whose spatial distribution overlaps with cortical 5-HT1B and 5-HT2A receptor densities. According to our biophysical model, these changes arise from the opposing effects of increased vasodilatory signal decay and reduced autoregulatory feedback on blood inflow. Our results also suggest that vascular effects may compromise the sensitivity of fMRI to detect psychedelic-induced alterations in inter-areal FC, rather than introducing spurious effects attributable solely to the vasoactive properties of these compounds. An optimistic interpretation is that the current literature may be overly conservative, underestimating the extent of FC alterations induced by psychedelics; however, further evidence is required to support this conclusion. Clarifying the magnitude of these confounds and their underlying mechanisms will be an important step toward a more complete understanding of psychedelic action, with far-reaching implications for basic neuroscience and clinical applications.

## Methods

### fMRI datasets

The analyses were performed on two previously published datasets of resting state fMRI data acquired under acute administration of LSD (Carhart-Harris et al., 2016) and DMT (Timmermann et al., 2023) in randomized, double-blind (LSD) or single-blind (DMT), within-subject designs. All participants provided written informed consent prior to their participation. Studies were approved by the National Research Ethics Service committee London-West London (LSD) and London—Brent (DMT), and research was conducted in accordance with the revised declaration of Helsinki, the International Committee on Harmonization Good Clinical Practice guidelines, and National Health Service Research Governance Framework. Imperial College London sponsored the research, which was conducted under a Home Office license for research with schedule 1 drugs

For LSD, 20 healthy participants were scanned during 14 minutes in two different days at least 2 weeks apart during eye-closed, task-free resting state. Scans were obtained after 135 minutes after infusion of LSD (75 μg) or an inactive placebo (saline). For DMT, 20 healthy participants were with simultaneous EEG and fMRI from 8 min before until 20 min after the intravenous injection of DMT (20 mg) or an inactive placebo (saline). Scans were obtained during eye-closed, task-free resting state. Further details on the experimental procedures, including drug administration and MRI data acquisition and preprocessing, are provided in the original publications (Carhart-Harris et al., 2016; Timmermann et al., 2023) and in the supplementary materials, sections “Experimental procedure (LSD)” and “Experimental procedure (DMT)”.

After data preprocessing, the datasets comprised 15 participants for the LSD experiment (four female, mean age 30.5 ± 8.0 years) and 17 participants for the DMT experiment (5 females, mean age 39.6 ± 9.6 years). Although the original datasets contained 217 time samples for the LSD experiment and 747 (drug condition) or 803 (placebo condition) time samples for the DMT experiment, all analyses were performed using temporal segments of equal length, to avoid biases due to time series duration. For the LSD dataset, the first 200 time points were analyzed. For the DMT dataset, a 300-volume segment corresponding to time points 200–500 was selected, as this interval encompasses the period of maximal subjective and physiological drug effects reported in the original study (Timmermann et al., 2023).

After preprocessing, BOLD time series were averaged at 90 cortical and subcortical regions of interest (ROIs) defined according to the Automated Anatomical Labeling (AAL) atlas, excluding the cerebellum (Tzourio-Mazoyer et al., 2022). For each of these time series, resting state HRFs were estimated according to the procedure explained in the following section.

### HRF estimation and deconvolution

Resting state hemodynamic response functions (HRFs) were estimated from the preprocessed BOLD time series using the rsHRF toolbox (Wu et al., 2013; Wu et al., 2021). The relationship between the underlying neural signal and these spontaneous events was modeled using the following linear time invariant (LTI) system:

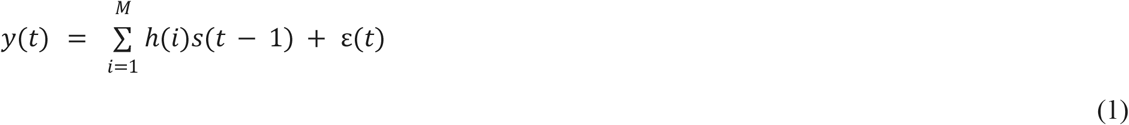

where *h*(*t*) represents the resting state HRF, *s*(*t*) is the spontaneous neural activity time series, and ε(*t*) is normally distributed measurement noise which is uncorrelated with *s*(*t*). The error term ε(*t*) has mean 0 and standard deviation of σ^2^*V*, with σ^2^ equal to the error variance and *V* given by the error correlation matrix. The length of the estimated HRF is given by parameter *M*.

To estimate the resting state HRF, *h*(*t*) is obtained using Eq. 1 by approximating *s*(*t*) with a point process ŝ(t), constructed from a set of delta functions that are centered at the onset of spontaneous BOLD events, identified as local maxima exceeding a threshold of one standard deviation above the mean signal (Tagliazucchi et al., 2012). This assumes that large amplitude excursions of the hemodynamic signals are driven by the spontaneous neural events encoded in ŝ (t). Because the BOLD signal lags behind the neural activation, the point-process is time-shifted by parameter τ, i.e.

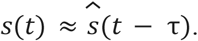

The optimal lag τ and the HRF *h*(*t*) are simultaneously obtained by minimizing the least-square error of the following optimization problem:

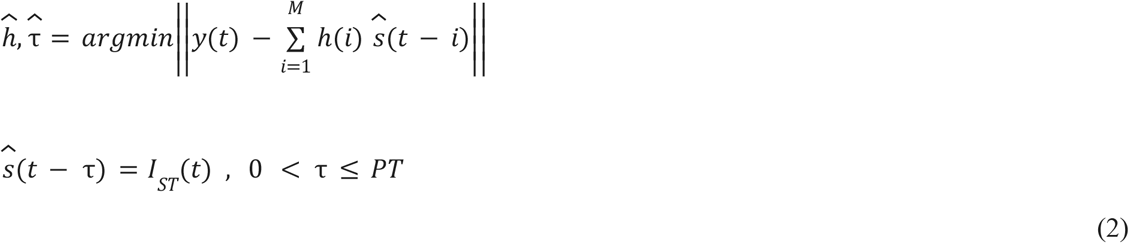

where ||·|| is the *L*_2_ norm, *ST* is the set of times where the BOLD signal has a local maxima and exceeds a threshold λ (1 s.d.), *PT* represents an upper bound to the hemodynamic delay (24 s), and *I*_*ST*_(*t*) is the indicator function for set *ST*, i.e. *I*_*ST*_ (*t*) = 1 if *t* ∈ *ST* and *I*_*ST*_ (*t*) = 0.

Finally, the LTI in Eq. 1 can be solved by implementing a Wiener deconvolution algorithm with the convolution kernel given by the HRF obtained from Eq. 2, ĥ (*t*). This operation resulted in the estimate of the neural activity time series *s*(*t*).

Three HRF shape parameters were extracted from the estimated HRFs ĥ (*t*): response height (RH), time to peak (TTP), and full width at half maximum (FWHM), which are taken as proxies for the hemodynamic response magnitude, latency, and duration (Lindquist et al., 2009). To improve the temporal resolution of these estimates, the HRFs were interpolated using cubic splines prior to parameter extraction. In cases where the ascending portion of the HRFs did not cross the half-maximum within the estimated time window, the initial segment of the response was linearly extrapolated from the first two interpolated samples to allow reliable estimation of the FWHM.

HRF parameters were first estimated for each AAL ROI and participant and subsequently averaged within six resting state networks (RSN), originally published by Beckmann and colleagues (Beckmann et al., 2005). Statistical comparisons (paired Student’s t-test) between drug and placebo conditions were then performed separately for each network and HRF parameter. The grouping by RSN was implemented to reduce the number of independent statistical comparisons between conditions and to facilitate the interpretation of the results.

### Receptor density maps

To examine whether psychedelic-induced changes in hemodynamic response latency were anatomically related to the cortical distribution of serotonin receptors, we compared regional changes in HRF time to peak (TTP) with receptor density maps derived from a previously published PET atlas (Hansen et al., 2013). We considered a subset of serotonin receptors (5-HT2A, 5-HT1B, and 5-HT1A) with established involvement in psychedelic pharmacology and vascular regulation. In particular, 5-HT2A receptor is the main target of psychedelic drugs and, together with receptor 5-HT1B, has been implicated with the vasoconstrictive effect of psychedelics, while receptor 5-HT1A has been primarily associated with vasodilation (Berger et al., 2009).

For each drug separately, regional drug–placebo differences in TTP were quantified using paired-tests across participants, yielding a t-value for each ROI. Pearson correlation coefficients were then computed between the regional t-value maps and each receptor density map. Statistical significance was assessed using permutation testing by randomly shuffling receptor density values across ROIs 50,000 times to generate null distributions of correlation coefficients. Empirical p-values were computed as the proportion of permutations yielding correlations equal to or greater than the observed value, and corrected for multiple comparisons using the false discovery rate (FDR) procedure with α =0.5.

### Functional connectivity and signal variance analyses

Functional connectivity (FC) analyses were performed separately on the original BOLD signals and on the corresponding deconvolved time series obtained with the rsHRF toolbox. For each participant, ROI time series were first averaged within the six resting state networks (RSNs) considered in this study, yielding one representative time series per network. Pairwise Pearson correlation coefficients were then computed between all RSN time series to obtain a 6 × 6 FC matrix for each participant and experimental condition.

To assess the effects of psychedelics on FC, drug–placebo difference matrices were computed separately for the original BOLD signals and for the deconvolved time series by subtracting the placebo FC matrix from the corresponding drug FC matrix for each participant. Paired t-tests were then performed across participants for each network pair, yielding statistical maps of drug-induced FC changes before and after deconvolution.

To evaluate the contribution of the estimated hemodynamic component to FC, comparisons between the original BOLD and deconvolved signals were performed separately for the drug and placebo conditions. For each participant, BOLD–deconvolved difference matrices were computed within each experimental condition, and paired t-tests were applied across participants for every network pair. Comparing the BOLD and deconvolved FC matrices within the same experimental condition allowed us to assess how the estimated hemodynamic component influenced functional connectivity under placebo and psychedelic conditions.

Signal variance was computed separately for the original BOLD and deconvolved time series. For each participant and experimental condition, the temporal variance was first calculated for each ROI and then averaged across all 90 regions, yielding a single mean variance value per participant.

Drug-induced changes in signal variance were assessed independently for the original BOLD and deconvolved signals by comparing the mean variance values obtained under the drug and placebo conditions using paired t-tests across participants.

### Dynamic mean field model

We employed a Dynamic Mean Field (DMF) network model to simulate spontaneous brain activity across the entire brain, with individual brain areas represented as nodes interconnected by white matter pathways (Deco et al., 2018). This model characterizes the activity within each brain region by simplifying the behavior of large ensembles of interconnected excitatory and inhibitory neurons into a set of dynamical equations, as established by Wong and Wang (Wong and Wang, 2006). In the DMF model, excitatory synaptic currents *I*^(*E*)^ are mediated by NMDA receptors, while inhibitory currents *I*^(*I*)^ are mediated by GABA_A_ receptors. Within each brain area *n*, excitatory (E) and inhibitory (I) neuronal pools are reciprocally connected, whereas inter-area coupling between two areas *n* and *p* only occurs at the excitatory-to-excitatory level, and is scaled by the structural connectivity matrix *C*_*np*_ and a global scaling parameter *G*.

The dynamics of the DMF model are described by the following system of coupled differential equations:

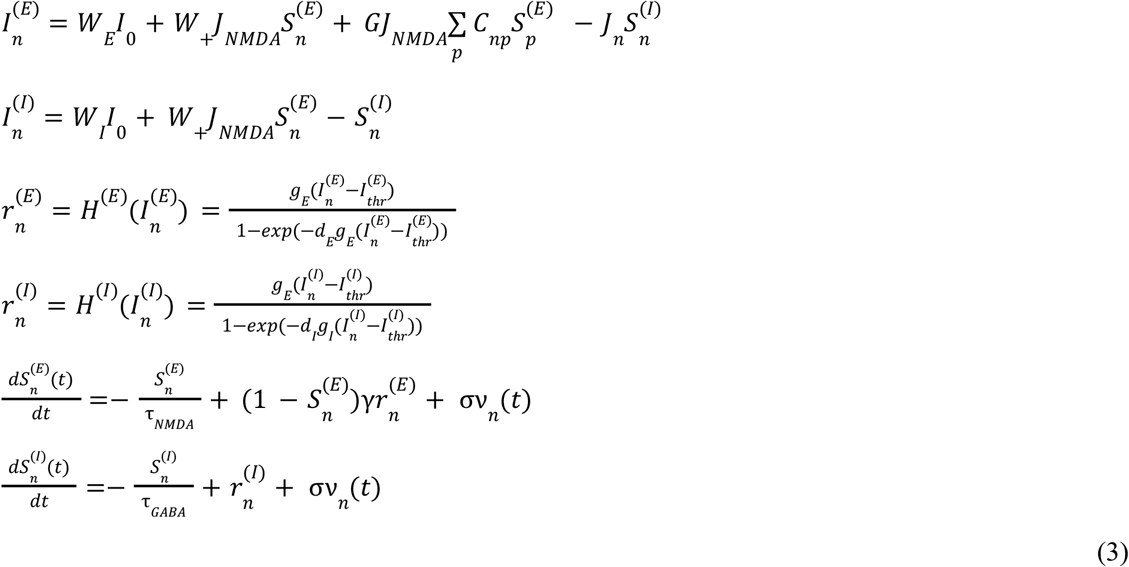

In each brain area *n*, the total input current for E and I pools is denoted by 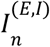 (in nA), while their firing rate is represented by 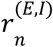 (in Hz). The neuronal response functions, denoted by *H*^(*E,I*)^, transform the total input currents received by the E and I pools into 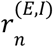 according to the input-output function described by Abbott and Chance (Abbot and Chance, 2005). Gain factors *g*_*E*_ = 310 nC^-1^ and *g*_*I*_ = 615 nC^-1^ determine the slope of *H*. Additionally, the threshold currents 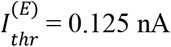 and 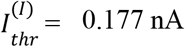 determine the points at which firing rates begin to increase linearly with input currents. Constants *d*_*E*_ = 0. 16 and *d*_*I*_ = 0. 087 shape the curvature of *H* around the threshold values.

The synaptic gating variable of excitatory pools, 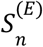, is regulated by NMDA receptors, is characterized by a decay time constant of τ_*NMDA*_ =0.1 s, while the synaptic gating in inhibitory pools is influenced by GABA receptors with a decay time constant of τ_*NMDA*_ =0.01 s. The overall effective external input is *I*_0_ = 0. 382 *nA*, with excitatory and inhibitory weights represented by *W*_*E*_ =1 and *W*_*I*_ =0.7, respectively. Additionally, *W*_+_ =1.4 is the weight of the recurrent excitation, while *J*_*NMDA*_ =0.15 nA globally weights all excitatory synaptic couplings. ν_*n*_ represents uncorrelated Gaussian noise with amplitude σ=0.01 nA. All parameter values are set according to previous work by Wong and Wang (Wong and Wang, 2006).

DMF model parameters were defined to replicate resting state conditions, such that each isolated node exhibits the characteristic noisy spontaneous activity with a low-frequency firing rate 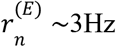 reported in electrophysiology experiments. The local feedback inhibitory control (FIC) parameter for region *n*, given by *J*_*n*_, was iteratively adjusted to maintain this firing rate (Deco et al., 2014).

Following the parcellation of structural and functional MRI data, we included N = 90 brain areas in our whole-brain network model. Brain are *n* receives excitatory input from all structurally connected areas, represented by *p*, into its excitatory pool. This input is weighted by the connectivity matrix, *C*_*np*_, derived from diffusion MRI data. Additionally, all inter-area excitatory-to-excitatory connections are uniformly scaled by a global coupling factor, denoted as *G*. This parameter was first adjusted at the subject-level to optimize to ensure that simulated activity closely matches the empirical resting state activity of each group, measured by the structural similarity index measure (SSIM) between simulated and group-average empirical FC matrices (Sanz Perl et al., 2021). Simulations were conducted across *G* values ranging from 1 to 2.5 with an increment of 0.01 and an integration step of 0.1 ms.

### Neurovascular coupling model

The simulated mean field activity was converted into a BOLD signal using a generalized hemodynamic model, proposed in Stephan et al. (2007). According to this model, the BOLD signal at area *n* is computed from the firing rate of excitatory pools, 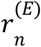. Increases in this firing rate are followed by increase of a vasodilatory signal, represented as *s*_*n*_, with spontaneous decay governed by time constant τ_*d*_. Blood inflow is denoted by *f*_*n*_ and responds proportionally to this signal, inducing a change in blood volume, *v*_*n*_, and deoxyhemoglobin content, *q*_*n*_. Changes in *f*_*n*_ exert regulatory feedback on *s*_*n*_ with a time constant given by τ_*f*_. The equation governing this biophysical model of the neurovascular coupling are as follows:

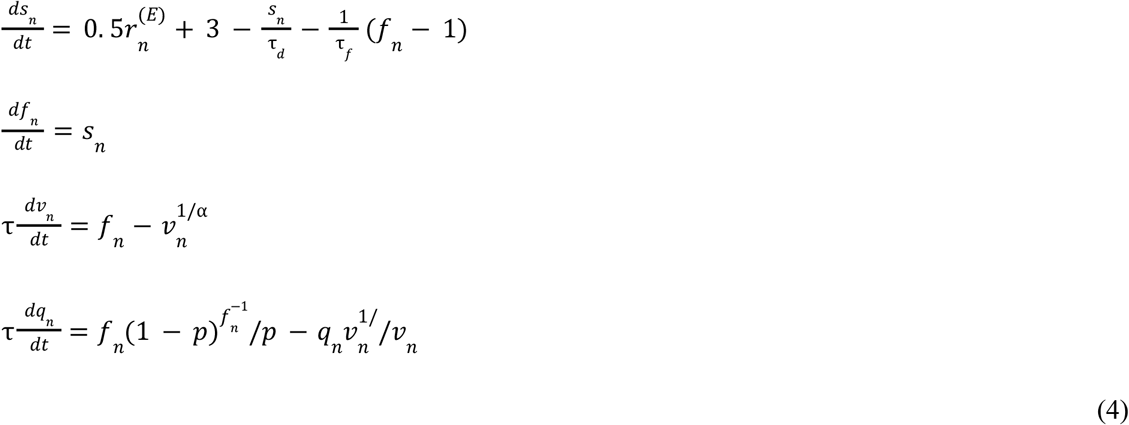

where *p* represents the resting oxygen extraction fraction, τ denotes a time constant for the changes in *v*_*n*_ and *q*_*n*_, and α represents vein resistance. The BOLD signal in each area is denoted by *B*_*n*_ is determined by the following nonlinear function of the blood volume and the deoxyhemoglobin content:

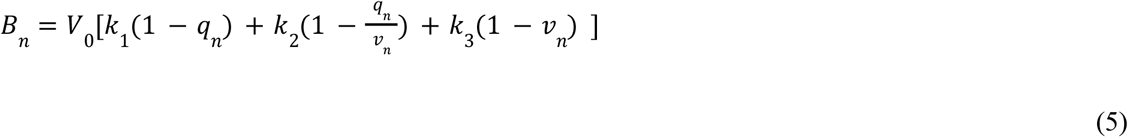

All model parameters were adopted from Stephan et al. (2007). To focus on the frequency range where resting state activity is most functionally relevant, both empirical and simulated BOLD signals were band-pass filtered between 0.01 and 0.08 Hz.

After an initial phase of parameter exploration, κ and γ were exhaustively varied between 0.04 and 0.6. For each group (LSD, DMT and corresponding PCB) and parameter combination, we used the rsHRF toolbox to estimate the time to peak (TTP) parameter of the simulated BOLD signals at each ROI, and the average TTP of the simulated BOLD signals was then compared with the empirical value.

### Statistical analyses

Comparisons between drug and placebo conditions, as well as between original BOLD and deconvolved signals where applicable, were assessed using paired two-tailed t-tests across participants. Correlations between regional changes in hemodynamic response parameters and serotonin receptor density maps were evaluated using Pearson correlation coefficients, and statistical significance was assessed using permutation testing with false discovery rate (FDR) correction for multiple comparisons. Unless otherwise specified, statistical significance was defined as (p < 0.05).

